# Do Support Vector Machines play a role in stratifying patient population based on cancer biomarkers?

**DOI:** 10.1101/2020.11.02.364612

**Authors:** Ben Lanza, Deepak Parashar

## Abstract

Biomarkers are known to be the key driver behind targeted cancer therapies by either stratifying the patients into risk categories or identifying patient subgroups most likely to benefit. However, the ability of a biomarker to stratify patients relies heavily on the type of clinical endpoint data being collected. Of particular interest is the scenario when the biomarker involved is a continuous one where the challenge is often to identify cut-offs or thresholds that would stratify the population according to the level of clinical outcome or treatment benefit. On the other hand, there are well-established Machine Learning (ML) methods such as the Support Vector Machines (SVM) that classify data, both linear as well as non-linear, into subgroups in an optimal way. SVMs have proven to be immensely useful in data-centric engineering and recently researchers have also sought its applications in healthcare. Despite their wide applicability, SVMs are not yet in the mainstream of toolkits to be utilised in observational clinical studies or in clinical trials. This research investigates the very role of SVMs in stratifying the patient population based on a continuous biomarker across a variety of datasets. Based on the mathematical framework underlying SVMs, we formulate and fit algorithms in the context of biomarker stratified cancer datasets to evaluate their merits. The analysis reveals their superior performance for certain data-types when compared to other ML methods suggesting that SVMs may have the potential to provide a robust yet simplistic solution to stratify real cancer patients based on continuous biomarkers, and hence accelerate the identification of subgroups for improved clinical outcomes or guide targeted cancer therapies.

## 1 Introduction

Cancer is a worldwide public health issue that affects millions of people every year. In 2018 there were 17 million newly documented cases of cancer globally (8.8 million in men and 8.2 million in women), leading to 9.6 million deaths [1]. Cancer is a vastly heterogeneous disease, with over 100 different types of cancer currently identified in humans [2]; the most common types of cancer are lung, female breast, bowel and prostate, these four types account for more than 40% of all new cancer cases [1]. The number of treatment options for many different types of cancer is growing; classical methods such as surgery, chemotherapy and radiotherapy are being continually used and refined, and novel targeted and immunotherapies are being created to tackle different cancers. Moreover, academic institutions and the pharmaceutical industry are providing more and more exciting avenues of research to help fight cancer. A relatively new emergence in the area of patient care is personalised medicine, an approach that allows doctors to choose treatments that will maximise the benefit to the patient by obtaining a genetic understanding of their disease. A range of genetic tests indicate which treatments a patients tumour will respond to, and to what degree, allowing doctors to choose a particular treatment regime that will have the best outcome for the patient and sparing the patient from exposure to treatments that will have little or no effect on them. Personalised medicine is still at quite a young stage and is not yet part of the routine care that patients receive. New treatments are being tested in clinical trials with the goal of their eventual use in the area of personalised medicine, there are many treatments that have already been approved by the US Food and Drug Administration (FDA) that target specific cancer mutations and fall under the umbrella of targeted therapy [3].

### 1.1 Stratifying patient population based on cancer biomarkers

A central concept underlying personalised medicine is that of ‘stratification’ [4]. In the most intuitive case, the patient population is stratified into two subgroups based either on the patient characteristics (age, gender etc.), disease characteristics (stage of disease, size of tumour etc.) or with increasing popularity ‘biomarkers’. Simply put, a biomarker is an objective, quantifiable characteristic that acts as an indicator of a normal biological process, pathogenic process or a pharmacologic response to a therapeutic intervention [5]. Biomarkers can be molecular, physiologic or biochemical, they provide a measurement of a biologically produced change in the body that can then inform on disease progression or other health conditions. For example, genetic and epigenetic changes associated with a cancerous tumour can alter various molecules (nucleic acids/proteins/hormones), detecting changes in any of these can inform decisions on diagnosis and monitoring of cancer. Biomarker-based stratification enables substantial improvements in patient care by directing therapies to specific groups of patients that are most likely to benefit.

Biomarkers in oncology are often used for either prognosis or prediction of treatment benefit [6]. A prognostic biomarker informs on the progression of disease in the absence of any intervention, and should have strong and statistically significant association with the clinical outcome of interest such as progression-free survival, overall survival, or quality of life. Therefore, a stratification based on prognostic biomarker signature distinguishes between subgroup of patients with relatively good outcomes and those with relatively bad outcomes irrespective of the treatment. On the other hand, for predictive biomarkers, one splits the patient population into subgroups sharing differential treatment benefit based on the biomarker signature, and for this reason these are also known as treatment-selection or therapy-guiding signatures. In both cases, patients are stratified into subgroups based on prognostic or predictive biomarker signatures depending on whether investigators aim to improve clinical outcomes or guide therapies [7]. In this work we have focused on prognostic biomarker datasets as these can offer real value in scenarios where treatment options may be limited, or if patients need to categorised into high or low risk, or to use as a stratification variable in a randomised clinical trial. However, the methods are applicable generically to all biomarker based stratifications.

Further, a biomarker can either be qualitative or quantitative: qualitative biomarkers are either present or not, for example a mutation in the p53 gene (associated with a variety of cancers [8]) can be measured as present or not. A quantitative biomarker however can be measured on a continuous scale, taking any number within some range, with this range being dependent on the value being measured. For example, many blood serum values are recorded as concentrations in the blood. An important feature often associated with quantitative biomarkers is the notion of a “cut-off” value, for which a value above (or below, depending on context) represents an extreme or abnormal case; this is directly associated with stratification as patients with biomarker values above a certain cut-off value may hypothetically have better clinical outcomes, and so this biomarker would define the subgroup of interest in this case.

### 1.2 Machine Learning methods for stratification

As eluded to above, in order for a quantitative biomarker to have practical applicability in clinical decision making concerning the prognosis of disease or treatment of patients, there is often an associated cut-off value which is used to dichotomise the patient population. Depending on the role of the biomarker, this cut-off value may flag whether a particular disease is present, define high/low risk patients or identify patient subgroups that are more likely to benefit from treatment more than others. Often this cut-off value can be derived from clinical knowledge, an example of this is blood pressure and its associated risks. Hypertension is defined as any blood pressure higher than 140/90mmHg [9], this increase in blood pressure puts extra strain on the blood vessels and heart and is associated with various life-threatening conditions such as heart attacks, strokes and aortic aneurysms. However, this cut-off value is not often known with precision and needs to be defined so that the biomarker stratifies the population in an optimal fashion. Determining this cut-off value can be a challenging task as the guidance from clinical knowledge is usually limited and data resources can be lacking. Furthermore, many downstream activities associated with the biomarker will be directly dependent on this choice of cut-off, therefore it has the potential to impact things like company expenditure, time spent on projects and the speed at which patients receive new treatments. Thus, cut-off choice is heavily scrutinised and so care needs to be taken to ensure the optimal choice is being made.

Machine learning methods are an attractive option in the area of stratification [10], there exist multiple supervised learning techniques which are used to classify data into groups in an optimal way. Machine learning (ML) is seen as a subset of Artificial Intelligence (AI), its primary aim is to provide a system with the ability to learn and improve automatically without the need for explicit human input. ML has a wide range of applications in engineering, physics, mathematics, health care, finance and practically any area that that is data-centric. ML methods can be broadly categorised into two sets; supervised and unsupervised learning. Supervised learning is the more common of the two, as the name suggests the machine “learns” in the presence of a “teacher”; this is accomplished by giving the machine training input data which is correctly labelled, so it can learn what a successful output is. The machine can then apply what it has learnt using the training data to a novel situation and arrive at a correct classification. An example of this is the classification of fruit: suppose that you have data on the size, shape and colour of certain known fruits:

| **Fruit** | **Size** | **Colour** | **Shape** |
| --- | --- | --- | --- |
| Apple | Big | Red | Round with a dent in the top |
| Banana | Big | Yellow | Long, cylindrical |
| Orange | Big | Orange | Round |
| Strawberry | Small | Red | Heart shaped/round |
| Grape | Small | Green | Oval |

Then, by using this training data to learn, the machine will then be able to make an educated guess that a novel fruit that is big, red and round with a dent in the top is an apple. Unsupervised learning is less commonly used but still has many useful applications. In this case, the training input data is unlabelled and the goal of the machine is to uncover an underlying structure or distribution from the data. Using a similar example as above, an unsupervised machine would be expected to form clusters of fruit based on their features i.e. separate the fruits based on their size, colour and shape. Therefore, it will be able to identify a group of big, orange, round fruits and separate this from a group of small, oval, green fruits, but will not have a name for what either of these fruits actually are.

Many supervised learning methods exist to carry out stratification, such as Decision Trees, Neural Networks and Naive Bayes methods [10], the work presented here focusses on the use Support Vector Machines (SVMs) [11]. SVMs are one of the most recent ML techniques and have shown applicability to a variety of real-world problems. Since being proposed by Vapnik and Lerner in 1963 [12] and formalised into the method known today in 1995 (by Vapnik and Cortes) [11], SVMs have been applied to text categorisation [13], tissue classification [14], gene function prediction [15], handwritten digit recognition [16] and facial recognition [17]. SVMs continue to be applied in biomedicine and healthcare, with researchers utilising them for cancer classification, biomarker discovery and drug discovery [18]. In the majority of cases, SVMs outperform other classification methods, it has been demonstrated that they achieve better predictive accuracy than Decision Tress [19], Random Forests [20] and Neural Networks [21]; SVMs also offer a range of benefits which make them an attractive option for stratification problems.

Despite the wide applicability of SVMs in biomedicine and healthcare, little work has been done with regard to stratifying patient populations based on prognostic or predictive biomarkers for applicability in observational studies and clinical trials. In this work we address the abovementioned gap by examining the role of SVMs in stratifying patient populations based on continuous biomarkers. In Section 2, we review the mathematical framework underlying SVMs including non-linear datasets. Section 3 sets out details for implementation of SVMs for patient stratification using a variety of prognostic biomarker datasets. Results of the analyses are presented in Section 4 followed by Discussion in Section 5.

## 2 Methodology of Support Vector Machines

As discussed, SVMs are a type of supervised machine learning method which are used to solve classification problems. Given data similar to that shown in Figure 1, a SVM should be able to classify the data points appropriately. Then, given a novel point with observed values for ‘example variable 1’ and ‘example variable 2’, will be able to classify it as ‘class 1’ or ‘class 2’.

**Figure (1).**
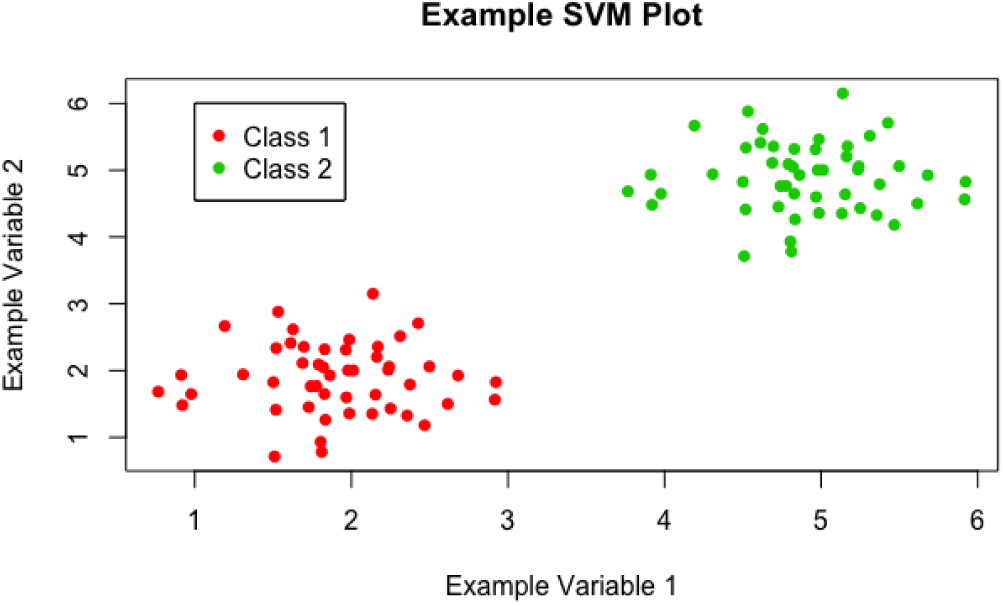
A scatter plot of example data, showing a classification problem to which SVMs can be applied

SVMs achieve this by constructing an ‘optimal hyperplane’ to split the given data; many hyperplanes have the ability to split the data, however the ‘optimal’ hyperplane is the one that maximises the margin, or distance, between the two classes of data (see Figure 2). Maximising this margin lends confidence to the method as novel data will be classified as accurately as possible. Here, a hyperplane is a decision boundary that classifies the data, depending on which side of this line a data point falls will define its classification. The dimensionality of the hyperplane is directly dependent on the number of input variables: in Figure 2, the number of input variables is 2, so the hyperplane is just a line; if there were 3 input variables, then the hyperplane would be a plane; for higher dimensions it becomes too hard to visualise, but if there were *N* input variables, the hyperplane would have *N* − 1 dimensions.

**Figure (2).**
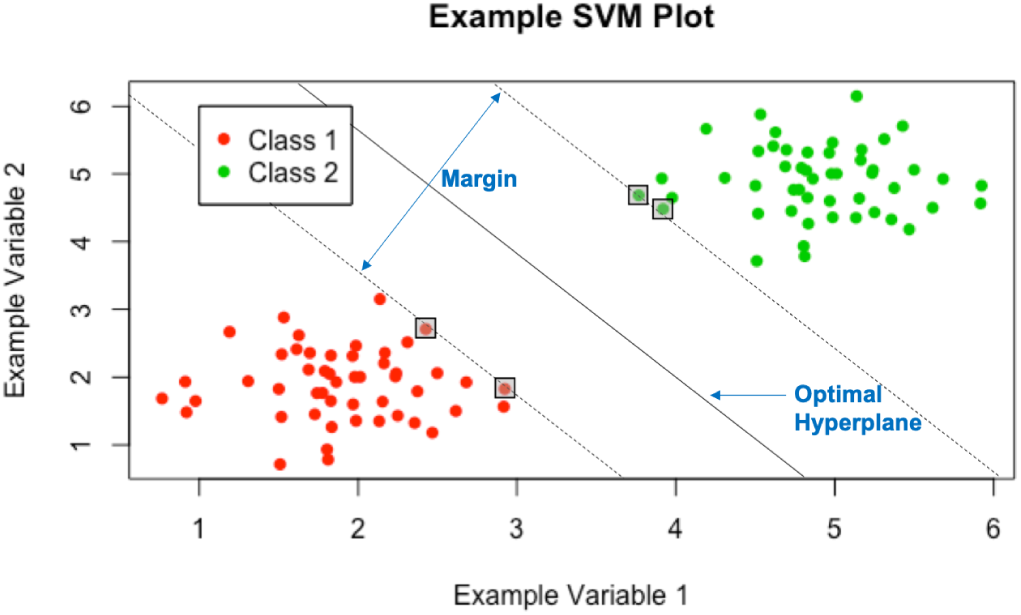
The optimal hyperplane yielded by the SVMs.

‘Support Vectors’ are the data points which lie closest to the optimal hyperplane and lie directly on the edge of the margin (and hence give the nomenclature Support Vector Machine), these points are denoted on Figure 2 in boxes. The support vectors are the most difficult data points to classify and directly influence the location of the optimal hyperplane; in fact they completely specify the mathematical form of the decision surface, which shall be seen. Now, the mathematical framework underlying SVMs will be explored, this work is based on the paper by Vapnik and Cortes [11].

### 2.1 Linearly separable case

Begin by looking at the simplest case, in which the data is linearly separable i.e. an optimal hyperplane can be constructed with associated margin, with no data points encroaching on the margin or falling on the incorrect side of the hyperplane (much like in Figure 1). This is rarely the case with real examples, but the mathematics of this base case can be extended to incorporate more difficult scenarios which will be explored in the following sections.

As with most classification techniques, there are associated **inputs** and **outputs**. In the case of SVMs, these are:

#### Inputs

A set of training pair samples (***x***_*i*_,*y*_*i*_), where ***x***_*i*_ ∈ ℝ^*n*^, *i* = 1, …, *l* is a feature vector of size n that contains all of the input variables for *l* different data points and *y*_*i*_ ∈ {−1, 1} is the classification for these *l* data points and takes the value either 1 or −1.

#### Outputs

A set of weights ***w*** ∈ ℝ^*n*^ and bias *b ∈* ℝ whose linear combination predicts the classification for a novel input vector ***u***. Non-zero weights are defined by the support vectors and hence there are a relatively small number of them, which is attractive computationally.

Some simple definitions are required before proceeding. Define the hyperplanes *H*_1_, *H*_2_ and *H*_0_ (see Figure 3) such that ***w***^*T*^ ***x***_*i*_ + *b ≥* 1 for *y*_*i*_ = 1 (the green portion of points on Figure 3) and ***w***^*T*^ ***x***_*i*_ + *b ≤* −1 for *y*_*i*_ = −1 (the red portion of points). NOTE: here ***w***^*T*^ ***x***_*i*_ denotes the inner product or scalar multiplication of the two vectors, also represented as ***w* · *x***_*i*_. Then *H*_1_ and *H*_2_ are the hyperplanes that define the edges of the margin and hence the support vectors lie on them:

**Figure (3).**
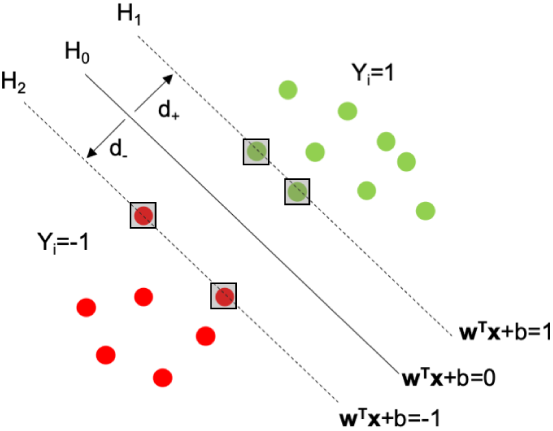
A plot to aid in defining the hyperplanes *H*_0_, *H*_1_, *H*_2_ and the associated margin.

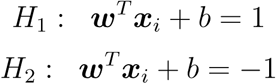

Then the decision surface, i.e. the optimal hyperplane, is represented by *H*_0_ and is the median between *H*_1_ and *H*_2_, it takes the form:

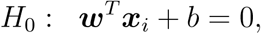

where ***w*** is the vector of weights, ***x***_*i*_ is an input feature vector and *b* is the bias. The margin of separation is the distance between *H*_1_ and *H*_2_, and is calculated as *d* = *d*_+_ + *d*_−_. The optimal hyperplane is the hyperplane that **maximises** this margin *d*.

The problem that needs to be formulated is to maximise this margin subject to the constraint that there are no data points within the margin. The size of the margin can be stated formally using the general equation for the distance from a point (*x*_0_, *y*_0_) to a line *Ax* + *By* + *c* = 0:

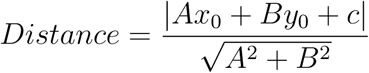

In this case, the perpendicular distance from a point on *H*_0_ to the hyperplane *H*_1_ is then:

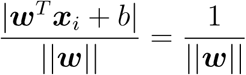

Doubling this to obtain the size of the margin gives 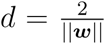. So, in order to maximise the margin 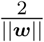, one needs to minimise ‖***w***‖.

The constraint of there being no data points can be written formally as ***w***^*T*^ ***x***_*i*_ + *b ≥* 1 for *y*_*i*_ = 1 and ***w***^*T*^ ***x***_*i*_ + *b ≤* −1 for *y*_*i*_ = −1; combining these for ease of notation gives *y*_*i*_(***w***^*T*^ ***x***_*i*_ + *b*) *≥* 1.

Therefore, the problem can be written formally as:

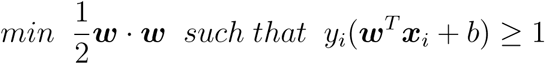

NOTE: ‖***w***‖ has been exchanged for 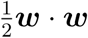 in order for standard optimisation techniques to be used. The same optimisation is still achieved.

The problem is now represented as a constrained, quadratic optimisation problem that can be solved using the Lagrangian multiplier method. The Lagrangian multiplier method is a technique for finding the local maxima or minima of a function subject to equality constraints. For the general case with the goal of maximising some *f* (*x*) subject to *g*(*x*) = 0, the Lagrangian is:

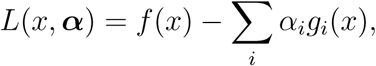

with the constraint that the partial derivates with respect to both *x* and ***α*** are equal to 0:

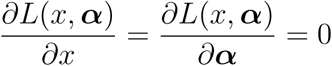

where the vector ***α*** contains the Lagrange multipliers, *α*_*i*_. Formulating the problem in this way ensures that the original problem is solved: forcing 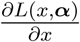 to be equal to 0 recovers the ‘maximal’/’minimal’ constraint, ensuring that the minimal solution is found; forcing 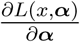 to be equal to 0 recovers the constraint of *g*(*x*) = 0, ensuring that all additional constraints are met. In the current case, 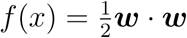 and *g*(*x*) = *y*_*i*_(***w**^T^**x**_i_* + *b*) − 1, so the Lagrangian is:

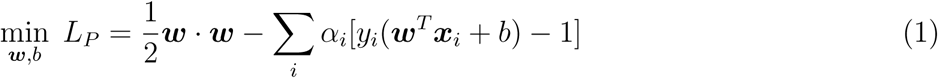

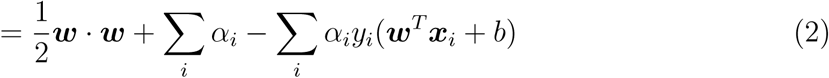

Then, by taking partial derivatives with respect to ***w*** and *b* as required:

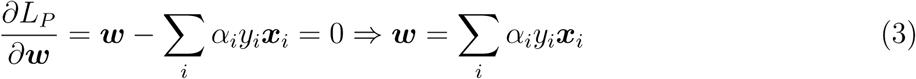

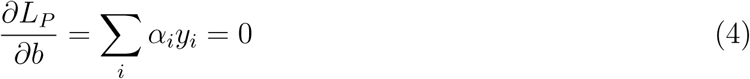

By observing equation 3, it is clear that the answer for the form of ***w*** has been obtained: ***w*** can be written as a linear combination of all *α*_*i*_, *y*_*i*_ and ***x***_*i*_; so all that remains is to find all *α*_*i*_. Most of these *α*_*i*_ will be zero, the non-zero *α*_*i*_ terms correspond to the support vectors.

The formulation of equation 2 is known as the **Primal** form of the optimisation problem. Instead of solving this, the **Dual** form of the problem will be solved. In the Primal form of the problem, one is minimising *L*_*P*_ (note the subscript P to represent the <u>P</u>rimal) over ***w*** and *b* with respect to constraints in terms of *α*_*i*_. In the Dual of the problem, one can maximise over the *α*_*i*_ (the Dual variable) subject to previously obtained constraints in terms of ***w*** and *b*; namely equations 3 and 4. Furthermore, by substituting the values obtained for these constraints into equation 2, the dependencies on ***w*** and *b* can be removed and the final formulation is in terms of *α*_*i*_ only.

The Dual of the problem is therefore:

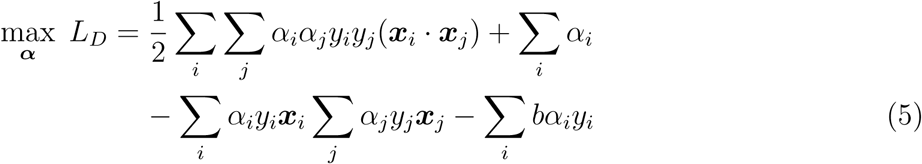

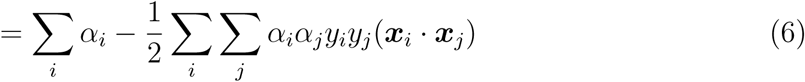

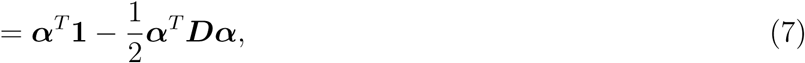

subject to the constraints ∑_*i*_ *α*_*i*_*y*_*i*_ = 0 and *α*_*i*_ *≥* 0, *i* = 1, …, *l*. Where **1** is vector of ones of length *l* i.e. **1** = (1, …, 1) and ***D*** is a square, symmetric matrix such that *D*_*ij*_ = *y*_*i*_*y*_*j*_(***x***_*i*_***x***_*j*_). Equation 7 is the same formulation of the problem obtained in 6, just represented in vector notation. This final form of the optimisation problem is easy to solve using existing quadratic programming methods, meaning that the *α*_*i*_ values can be obtained and ***w*** can be calculated. Furthermore, the value for the bias, *b*, can also be obtained by calculating *b* = *y*_*i*_ − ***w***^*T*^ ***x***_*i*_ for each support vector and then averaging over all results.

Finally, with values obtained for ***w*** and *b*, given a novel feature vector ***u***, one can obtain its predicted classification based on its observed input variable values by observing on which side of the decision surface it lies; this is achieved by taking the sign (is it positive or negative) of the decision function:

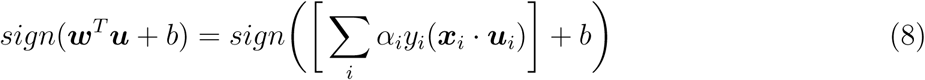

### 2.2 Non-linearly separable cases

The linear case is rarely seen in the real world, there is often no clear point at which to separate two classes, as can be seen in Figure 4. There are two techniques, often used in conjunction, that can be applied to deal with this issue.

**Figure (4).**
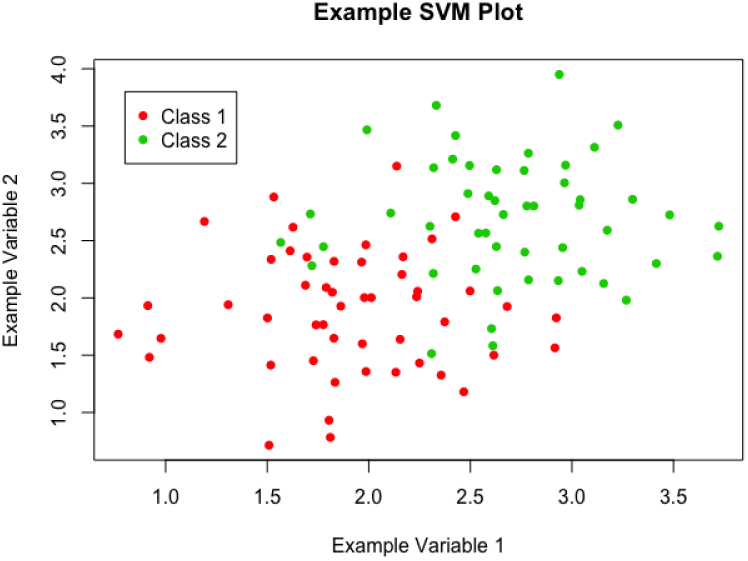
A scatter plot of example data, showing a more realistic scenario, in which the data is not linearly separable

#### Soft-Margin

The idea behind the ‘soft-margin’ approach is to allow some mistakes to be made when the algorithm is learning to classify the data. This is achieved by the introduction of new slack variables *ξ*_*i*_, which soften constraints. In the original formulation of the problem, the constraint that no points could fall within the margin was represented as *y*_*i*_(***w***^*T*^ ***x***_*i*_ + *b*) *≥* 1; following the introduction of new slack variables, this becomes:

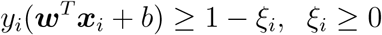

This implies that the new functional margin is allowed to be less than 1 and the problem also makes use of a cost parameter C (introduced below) that penalises in two cases:

**Case 1**: Points that fall on the correct side of the decision surface but within the margin i.e. 0 *< ξ*_*i*_ *≤* 1.

**Case 2**: Points that fall on the incorrect side of the decision surface and hence have incorrect classification, according to the decision function i.e. *ξ*_*i*_ *>* 1.

Thus, there is a built in preference for margins that classify the data correctly, but the constraints have been softened to allow for non-separable data, with an additional penalty that is proportional to the amount of data that is misclassified. The problem is therefore represented as:

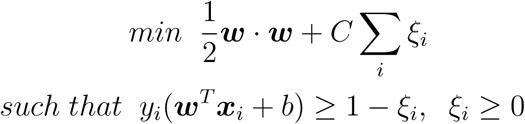

In practice, the optimal value for *C* is determined by cross-validation; this involves validating a number of separate models with different values for C and observing which performs the best. The mathematics follows in a similar manner to the linearly separable case, a Lagrangian is initially constructed:

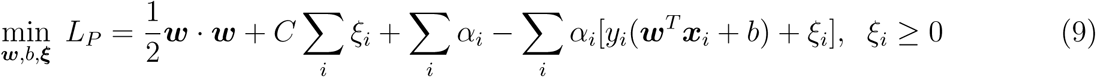

Again, equation 9 represents the Primal of the optimisation problem, the Dual will be solved instead. The Dual is constructed in the same fashion as in the linearly separable case:

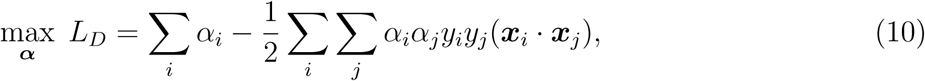

subject to the constraints 0 *≤ α*_*i*_ *≤ C* and ∑_*i*_ *α*_*i*_*y*_*i*_ = 0. The box constraint on the *α*_*i*_ is necessary to ensure that the Lagrangian function is bounded i.e. that the cost cannot be driven to −*∞*. This occurs when *C* − *α*_*i*_ *<* 0, *ξ*_*i*_ approaches *∞* and so ∑_*i*_(*C* − *α*_*i*_)*ξ*_*i*_ goes to − *∞*. The box constraint works to prevent this; the problem is the same as in the linearly separable case except for the introduction of this box constraint. The problem then follows in the same manner as before, the *α*_*i*_ are obtained by solving equation 10, allowing one to obtain an expression for ***w*** and *b* and hence the decision function.

#### Transformations to a Higher Dimensional Space (Kernel Trick)

All methods so far have depended on the inner products between sets of input data points i.e. ***x***_*i*_ *·* ***x***_*j*_ or equivalently 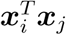. As a result of this feature, these methods can be applied to nonlinear situations very easily. The aim here is to gain linear separability by transforming the data to a higher dimensional space, or applying some function to the original data to change its appearance, an example of this is given in Figure 5. The data in Figure 5a is clearly not linearly separable, but after applying a transformation, a plane can be drawn to separate the two classes in Figure 5b. Data is mapped to this higher dimensional space by applying a function to the whole dataset, for ease of notation call this function *φ*(), but this can represent any appropriate function.

**Figure (5).**
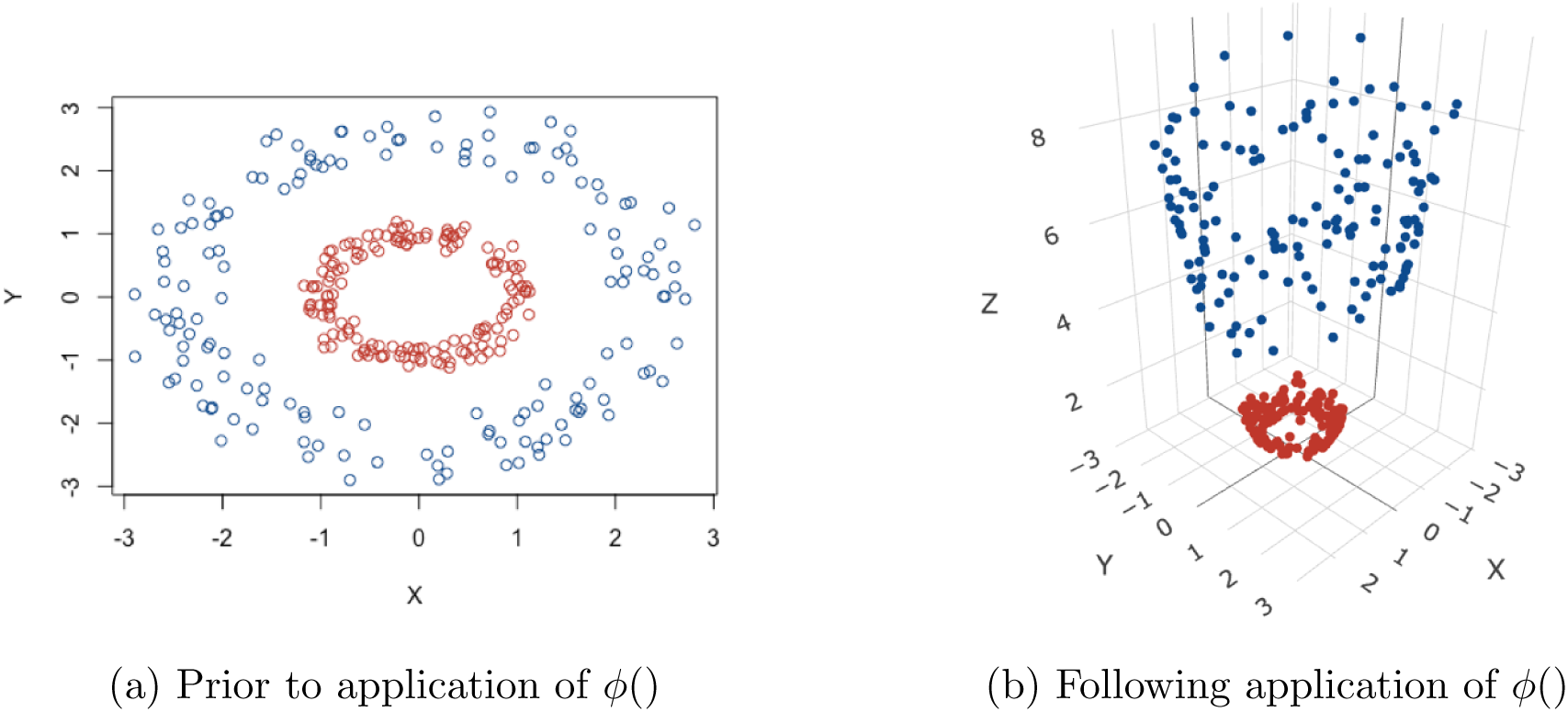
The ‘Kernel Trick’: Transforming data to higher dimensions to gain linear separability [22]

Following application of *ϕ*() to the data, the mathematics follows through similarly to the linearly separable case. The optimisation functions becomes:

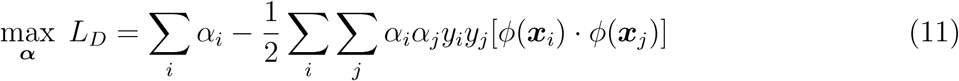

By looking at equation 11, it is clear that the solution is directly dependent upon calculating *ϕ*(***x***_*i*_) · *ϕ*(***x***_*j*_). This calculation can be very computationally expensive and time consuming or even impossible, and so it would be beneficial if this could be avoided. At this point, a Kernel function *K*(*x, y*) can be introduced to circumvent this issue. A Kernel function *K* is such that *K*(***x***_*i*_, ***x***_*j*_) *≡ ϕ*(***x***_*i*_) · *ϕ*(***x***_*j*_), meaning that *K*(***x***_*i*_, ***x***_*j*_) and *ϕ*(***x***_*i*_) · *ϕ*(***x***_*j*_) are equivalent for all pairs of data points ***x***_*i*_ and ***x***_*j*_; thus *K* essentially defines the inner product in the transformed space and allows one to obtain every value of *ϕ*(***x***_*i*_) · *ϕ*(***x***_*j*_) without needing to explicitly carry out a potentially complex calculation. Then, the inner product following transformation does not need to be computed at all, in fact one does not even need to know the form of *ϕ*() or even carry out the mapping to a higher dimensional space, as *K* gives all the information that is required.

The optimisation function then takes the form:

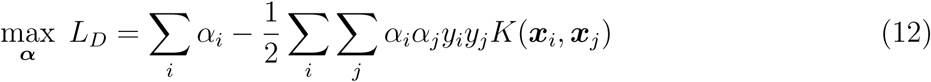

Calculation of the *α*_*i*_ and hence ***w*** and *b* follows as before. However, not all functions are viable to be used as Kernel functions; to be used as a Kernel function, a function *K*(*x, y*) needs to satisfy Mercer’s theorem [23]:

**Mercer’s Theorem**

A symmetric function *K*(*x, y*) can be expressed as an inner product

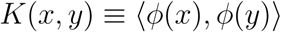

for some *ϕ if and only if K*(*x, y*) is positive semidefinite, i.e.

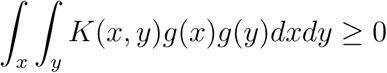

for any function g.

Some examples of widely used Kernel functions are given:

– **Polynomial**: (***x*** · ***y*** + *γ*)^*d*^
– **Radial Basis Function (RBF)**: 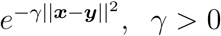
– **Sigmoid**: *tanh*(*κ****x*** · ***y*** − *δ*)

Note that different Kernel functions have different numbers of hyper-parameters, which needs to be accounted for during analysis.

### 2.3 Considerations when applying SVMs

SVMs are a powerful tool to be used for data classification problems, but results can be dependent on a number of choices when building the machine. These considerations will be explored in this section.

#### Data scaling

Before applying SVMs to a dataset, it is necessary that all input variables are continuous numbers and not categorical i.e. *x* = 1.47 instead of *x* =‘red’. Categorical attributes can be converted into a number of different variables which together account for its impact; an m-level categorical variable can be represented as m different continuous variables. Following this, all input variables need to be scaled to the same range; Hsu, Chang and Lin [24] recommend scaling all input variables to the range [-1,+1] or [0,1]. Scaling is a very important part of data pre processing as it stops input variables that cover a large range from dominating those that cover smaller ranges when applying the SVM algorithm. Furthermore, having all input variables on a smaller range avoids numerical problems when applying Kernel functions.

#### Choice of kernel function

As shown in section 2.2, there are 4 commonly used Kernel functions, the linear case (identity Kernel function), the radial basis function, the polynomial function and the sigmoid function. The choice of which to use is often data-dependent but in most cases, different Kernel functions are used in separate SVM models and the model that shows the best performance is used. Hsu, Chang and Lin [24] state that the RBF Kernel is a good first choice as it has many advantages over other Kernels and performs better in the majority of cases. Firstly, unlike the linear Kernel, it can handle non-linear data which makes it more widely applicable. Secondly, unlike more complex Kernels, the RBF Kernel only has two hyper parameters (the cost parameter C and *γ*); having a larger number of hyper-parameters to choose makes model selection more complex. Finally, the RBF Kernel has fewer numerical difficulties, the polynomial Kernel can sometimes diverge to *∞* when the *degree* parameter is large. However, the RBF Kernel is sometimes unsuitable, particularly in the case when the number of input feature variables is very large; in this case the linear Kernel is used.

#### Tuning of hyper-parameters: Cross-validation and grid search

Following the choice of appropriate Kernel function, hyper-parameters for this Kernel function must be chosen. Suppose the RBF Kernel has been selected, then the parameters *C* and *γ* must be chosen, the goal is to find the ‘best’ pair (*C, γ*) so that the SVM model can classify the data as accurately as possible. A common method when choosing parameters or training a machine learning technique is to split the data set into two parts, the first of which is used to ‘train’ the dataset and the second is considered unknown data and is used to validate the method; the accuracy of classification achieved on this validation portion of the dataset reflects the SVM’s performance on a set of novel data. An extension of this method is k-fold cross-validation: the training dataset is split into k equally sized subsets, the SVM is trained on a combination of k-1 subsets and then validated using the remaining subset. This is repeated until all subsets have been used as a validation set, so the same SVM model will be validated k times. This utilises the entirety of the dataset, and the accuracy of the SVM can be measured as the percentage of the data that has been correctly classified. K-fold cross-validation solves the issue of ‘overfitting’, an issue that arises when a SVM model is too closely fitted to the training dataset and so has little utility when classifying novel data.

A ‘grid search’ can then be carried out alongside cross-validation to find the optimal parameters, in this case *C* and *γ*. A number of pairs of (*C, γ*) are used in separate SVM models and the model that achieves the best cross-validation accuracy is then carried forward. Consider a simple example: try cost parameters *C* = (1, 2, 3) and *γ* = (0.5, 1), then the following pairs will be used in cross-validation and the best pair chosen for the final SVM model (*C, γ*) = (1, 0.5), (1, 1), (2, 0.5), (2, 1), (3, 0.5) and (3, 1). The different SVMs have different levels of complexity, in terms of hyper-parameter selection: the linear SVM only requires the cost value, the SVM with RBF Kernel requires the cost value and *γ* and the SVM with polynomial Kernel requires the cost value, *γ* and *d* (the degree).

## 3 Implementation of SVM for patient stratification

Having reviewed the SVM methodology in the preceding section, we now set out the algorithms necessary for stratifying the patient population into subgroups based on prognostic biomarker signatures. A brief overview of the datasets that were used to demonstrate classification using SVMs is given. A similar algorithm was used for data analysis in all cases and is outlined below (any major deviations from this algorithm for any datasets are detailed in their respective headings). All datasets used were either self simulated or publicly available as research and teaching resources.

### 3.1 Algorithm for application of SVM

All work presented here was done using R, the ‘e1071’ package [25] was used extensively when applying SVMs; the ‘e1071’ package is the R interface to the C++ implementation by Chih-Chung Chang and Chih-Jen Lin (‘libsvm’ [26]) and was created by David Meyer. The ‘dplyr’, ‘plotly’ and ‘caret’ packages were also extensively used. Prior to fitting any SVM models, simple data cleaning procedures were carried out. This was minimal in most cases because, as aforementioned, the datasets used were either self simulated or are publicly available to be used for teaching so they were in a usable condition to begin with. Simple preliminary analyses were also carried out on the data in order to ascertain the underlying distributions of biomarkers in each dataset and the proportion of patients that had the disease of interest (disease here is being used as a broad term to classify those patients that had the event under consideration in each dataset).

The respective dataset was then split into a training portion and a testing/validation portion; a 3:1 ratio was chosen for this split, so the training set contained 75% of the original data and the testing set 25%. 10-fold cross validation in conjunction with a grid-search was then carried on the training dataset. This was done to find the optimal hyper-parameters for each SVM with its respective Kernel function; the linear, RBF and polynomial Kernels were all explored, so this procedure was repeated three times in each dataset. The sigmoid Kernel was not used as it is only a valid Kernel function under certain conditions [27]. A soft margin approach was used alongside each Kernel, so the grid search was used to find the optimal cost parameter as well as hyper-parameters for each Kernel. The following sets of hyper-parameters were used when performing the grid search: cost value *C* = {2^−5^, 2^−4^, …, 2^4^, 2^5^}, *γ* = {0.25, 0.5, 1, 2, 4} and *d* = {1, 2, 3}.

After finding the optimal hyper-parameters in each case, all three models were then trained on the training data set and tested on the testing data set; scaling of the dataset was carried out prior to training the SVM models and was handled automatically by functions in the ‘e1071’ package. Testing here means predicting what classification the SVM would give to the ‘novel’ data, represented by the testing set. By comparing the predicted values to the observed classifications, a number of performance measures were calculated:

– 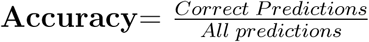, this is the proportion of the correct predictions given by the SVM
– 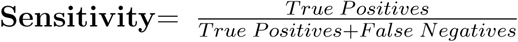, this represents the proportion of positive predictions that were correctly identified (a ‘positive’/ ‘negative’ depends on context, but a positive generally means that the disease is present).
– 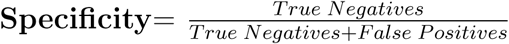, this represents the proportion of negative predictions that were classified correctly.

Additionally, the accuracy was compared to the No Information Rate (NIR) using an exact binomial test to obtain a formal comparison between the two, giving a one-sided P-value. The NIR is a naive method of classification, it is the proportion that would be classified correctly if every patient was classified as the majority class. For example, say that 70% of patients in a dataset had the disease of interest, if a classifier simply predicted that every patient was positive for the disease of interest, then it would achieve 70% accuracy; this is represented by the NIR. Therefore, the performance of the SVM can be compared to the simplest possible classifier by contrasting its performance with the NIR, this comparison is formalised using the exact binomial test. Moreover, the NIR can often give context to a high accuracy SVM classifier: a SVM may achieve 90% accuracy and therefore be a good candidate for further use, but if the NIR is actually 95%, then it is clear that this classifier does not actually have much utility in this case. Plots to explore how accurately the SVM model classified the data were then created, these displayed the cut off defined by the SVM with real classification values overlaid, so that performance could be assessed visually.

Finally, comparisons to other classification methods were carried out by comparing the performance measures as shown above. A logistic regression (LR) model and the k-Nearest Neighbours (kNN) algorithm were used for comparison as they are both popular choices for binary classification problems, such is the case here, and they are relatively simple to implement. The technical details of these methods are beyond the scope of this report, although both are briefly summarised here. A logistic regression model is a generalised linear model that uses a logistic function in order to model a binary outcome. The k-Nearest Neighbours algorithm is a non parametric supervised machine learning method with applications to classification and regression [28]. The general principle behind kNN is to find the *k* data points (*k* is predefined) that lie closest to a particular new data point and use the classification of these *k* points to predict the classification for the new data point; the Euclidean distance is the most commonly used distance measure when applying kNN algorithms, this was used here. Training and testing of the two alternative methods was carried out using the same approach to the SVM method; the coefficients of the LR model were obtained from the training portion using maximum likelihood estimation. No formal testing was carried out to compare SVMs to the two alternative methods, the performance measures were simply contrasted to informally investigate the performance of the various techniques.

#### Datasets for prognostic biomarkers

To demonstrate the applicability of SVMs for stratification based on prognostic cancer biomarkers, SVMs were applied to 5 datasets; 1 was self-simulated and 4 were publicly available. All datasets contained at least one continuous, prognostic biomarker measurement and a binary variable indicating whether the patient had the disease of interest. An overview of the datasets used is given below.

##### Self simulated Data

This dataset consisted of 200 observations of 4 ‘biomarker’ variables and a ‘disease’ flag. The ‘biomarker’ values were drawn from normal or exponential distributions (*B*1 *∼ N* (50, 10), *B*2 *∼ N* (200, 20), *B*3 *∼ exp*(0.1) and *B*4 *∼ exp*(0.01)). The outcome flag was defined using these biomarker values, any patient that had biomarker values in the top quartile for B1/B2/B3 simultaneously or B4 alone was considered as having the disease of interest. This dataset was used initially to practise implementation of SVM methodology.

##### Case study of prostate cancer

This dataset was made available by Etzioni et al. [29] and was downloaded from the Fred Hutch Diagnostic and Biomarkers Statistical (DABS) Centre website [30]. It contained 140 observations of 5 variables: a patient ID variable, a flag for whether prostate cancer was present, the patients age, a value for total prostate specific antigen (PSA) and a value for free PSA. For patients with multiple entries, the most up to date observation was used for analysis. The ratio of free to total PSA was then calculated as the measure of interest, PSA is the only widely used biomarker for diagnosis and prognosis of prostate cancer [31].

##### Pancreatic cancer biomarkers

This dataset was made available by Wieand et al. [32] and was downloaded from the Fred Hutch Diagnostic and Biomarkers Statistical (DABS) Centre website [30]. This dataset contained 141 observations of 3 variables: a flag for pancreatic cancer and two anonymised prognostic biomarker values. This dataset was included in analysis as a large skew was present in both biomarker values and how this affected SVM performance was of interest.

##### Gene expression array

This dataset was made available by Pepe et al. [33] and was downloaded from the Fred Hutch Diagnostic and Biomarkers Statistical (DABS) Center website [30]. This dataset contained 53 observations of 1,537 variables: a disease flag and 1,536 gene expression values, note that 12 gene variables were removed due to incomplete data, leaving 1,525 genes. It was included in analysis as it represented a ‘wide’ dataset and how this affected SVM performance was of interest.

##### Risk reclassification (simulated)

This dataset was made available by Pepe [34] and was downloaded from the Fred Hutch Diagnostic and Biomarkers Statistical (DABS) Center website [30]; this dataset was simulated. It consisted of 10,000 observations of 5 variables: a patient ID, a disease flag and 3 normally distributed biomarker values. It was included in analysis as it represented a ‘long’ dataset and how this affected SVM performance was of interest.

## 4 Results

### 4.1 Preliminary Analyses

Simple preliminary analyses were carried out prior to fitting SVMs to explore the biomarker distributions and proportion of patients with the disease of interest in each dataset.

#### Self simulated dataset

As this dataset was self simulated, the underlying biomarker distributions were already known, see section 3.1. Also detailed in section 3.1 is the method of ascertaining which observations were flagged for disease, using this method, 53 patients (26.5%) were flagged for disease and 147 (73.5%) were not.

#### Case study of prostate cancer

The initial structure of this dataset was longitudinal, it contained multiple assessments of total PSA and free PSA per patient over a given period of time. The disease status for each patient was the same at each visit for every patient, therefore, only the latest visit for each patient was taken forward for analysis so that there would be a unique observation per patient and it was thought that the most up to date value would be the most indicative of disease status.

The original data set contained 643 observations for 141 patients, the dataset used for analysis therefore consisted of 141 observations with one per patient. One patient was also dropped from analysis because they had a free PSA value of 100%, when all other values were less then 12%. The ratio of free PSA to total PSA is a commonly used [35] metric, therefore this was calculated for all patients and used as the sole feature variable when fitting the SVM. A histogram of the PSA ratio is given in Figure 6, the distribution was close to normal, with a left skew showing a larger number of lower ratio values. In this dataset, 70 (50%) of the 140 patients were flagged for disease.

**Figure (6).**
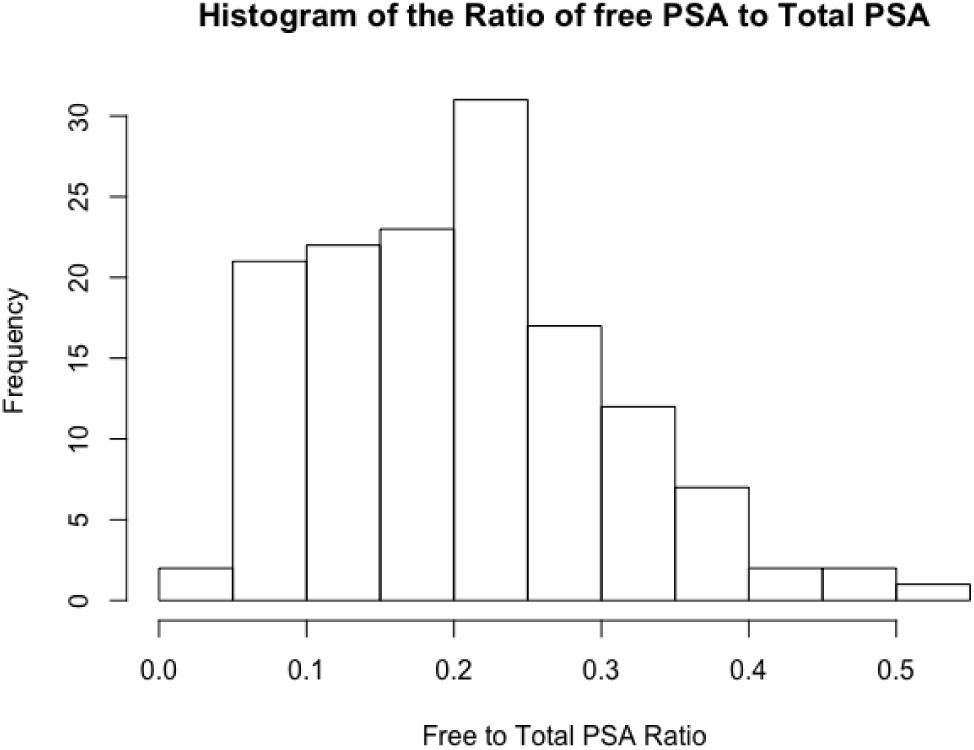
A histogram showing the distribution of the free to total PSA ratio among patients in the case study of prostate cancer dataset

#### Pancreatic cancer biomarkers

As discussed, this dataset was included to explore the impact of transformations on the SVM performance. The two anonymised variables y1 and y2 were extremely skewed, as can be seen in Figure 7a, the majority of data points were concentrated at lower values but there were numerous points at extreme values. After applying a log transformation to both y1 and y2, the plot of log(y1) vs log(y2) looked much better, with the data covering a sensible range and a visible distinction between the two classes (Figure 7b). In this dataset, 90 (63.8%) of the 141 patients were flagged for disease.

**Figure (7).**
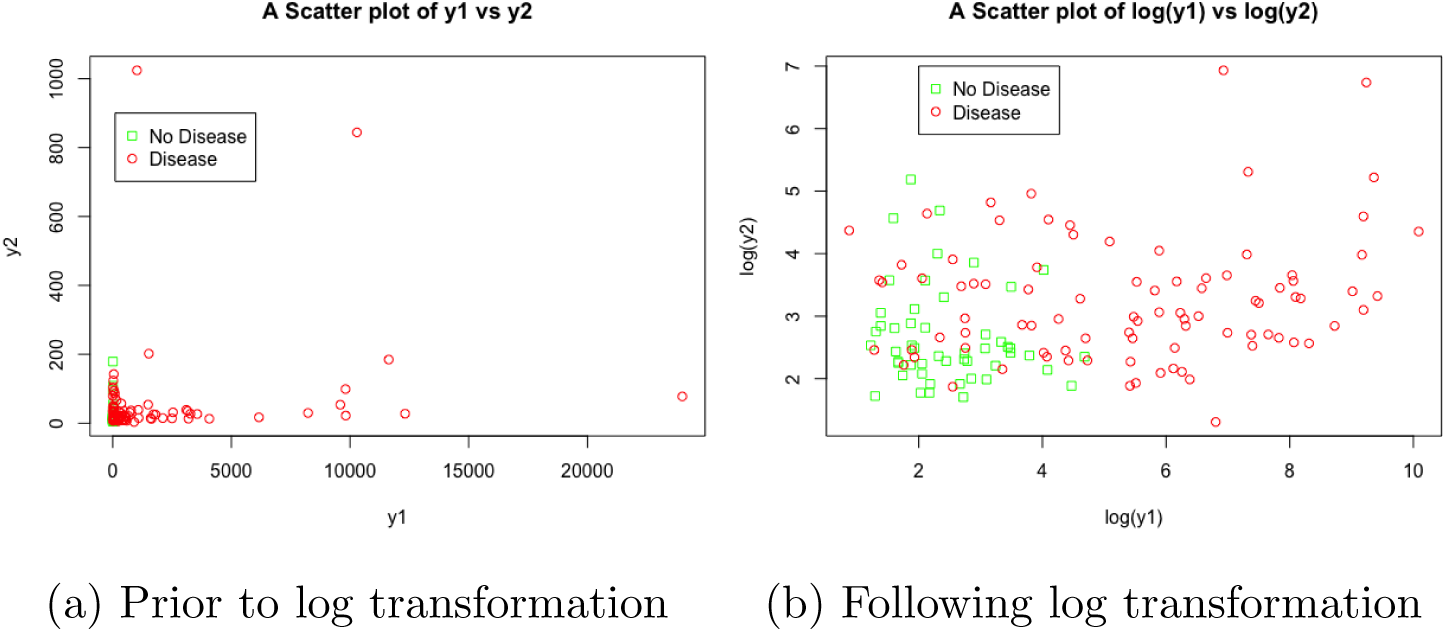
Scatter plots showing the relationship between y1, y2 and disease status, before and after transformation, in the pancreatic cancer biomarker dataset

#### Gene expression array

Due to this dataset containing 1,536 gene expression variables, no exploratory analyses were carried out to explore their distribution, this was because it was of more interest what impact on SVM performance including this large number of feature variables would have. However, it was discovered that 12 gene expression variables contained some missing values, so these were dropped from analysis and the final dataset contained 1,524 variables. Of the 53 patients included in analysis, 30 (56.6%) were flagged for disease.

#### Risk reclassification (simulated)

Histograms of the three simulated variables are given in Figure 8, it is clear that they were drawn from normal distributions centred around 0, with similar variances. A 3D scatter plot of all variables, split by disease status is also given in Figure 9, there is a clear distinction between the two classes; those with higher values of x, y and w seemed to have the disease of interest. Of the 10,000 patients simulated, 1017 (10.2%) were flagged for disease.

**Figure (8).**
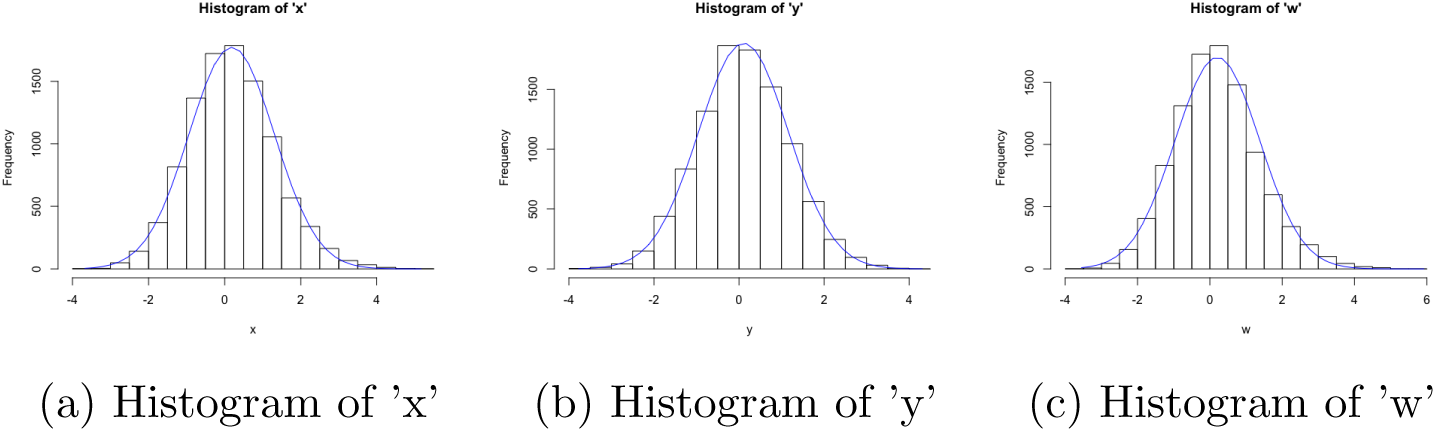
Histograms of all simulated variables in the risk reclassification dataset, with overlaid normal curves

**Figure (9).**
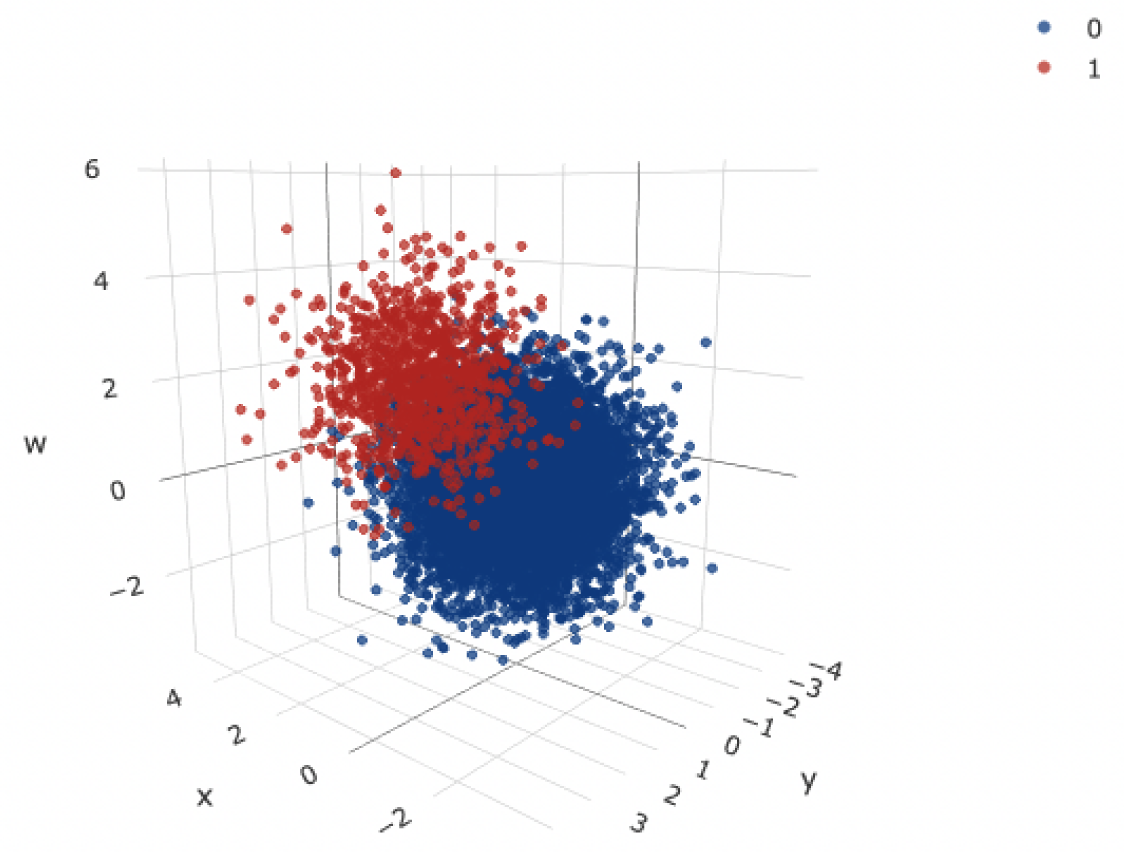
A 3D scatter plot of all simulated variables in the risk reclassification dataset, with disease status indicated (0-no disease, 1-disease)

### 4.2 Fitting SVMs

Table 1 gives the performance measures for SVM, logistic regression model and kNN algorithm used in each dataset. Details of model creation is given in each subsection prior to a comparison of final SVM model to the logistic regression model in each case.

**Table (1).** A table summarising the performance of all SVMs, logistic regression models and kNN algorithms. *-NIR is the No Information Rate in each dataset and the corresponding P-value was obtained by comparing the accuracy of each classifier to the NIR using an exact binomial test

| Method |  | Accuracy | Sensitivity | Specificity | NIR (P-value)* |
| --- | --- | --- | --- | --- | --- |
| Self-Simulated | SVM | 0.94 | 0.79 | 1.00 | 0.72(P<0.001) |
|  | Logistic | 0.94 | 0.86 | 0.97 | 0.72(P<0.001) |
|  | kNN | 0.90 | 0.71 | 0.97 | 0.72(P=0.002) |
| Case Study of Prostate Cancer | SVM | 0.72 | 0.79 | 0.65 | 0.53(P=0.014) |
|  | Logistic | 0.72 | 0.79 | 0.65 | 0.53(P=0.014) |
|  | kNN | 0.75 | 0.79 | 0.71 | 0.53(P=0.005) |
| Pancreatic Cancer Biomarkers | SVM (log) | 0.86 | 0.88 | 0.82 | 0.69(P=0.018) |
|  | SVM (no log) | 0.86 | 0.80 | 1.00 | 0.69(P=0.018) |
|  | Logistic | 0.81 | 0.88 | 0.64 | 0.69(P=0.099) |
|  | kNN | 0.78 | 0.84 | 0.64 | 0.69(P=0.184) |
| Gene Expression Array | SVM | 0.86 | 0.83 | 0.88 | 0.57(P=0.025) |
|  | Logistic | 0.43 | 0.17 | 0.63 | 0.57(P=0.911) |
|  | kNN | 0.93 | 1.00 | 0.88 | 0.57(P=0.005) |
| Risk Reclassification | SVM | 0.96 | 0.77 | 0.98 | 0.90(P<0.001) |
|  | Logistic | 0.96 | 0.74 | 0.98 | 0.90(P<0.001) |
|  | kNN | 0.96 | 0.78 | 0.98 | 0.90(P<0.001) |

#### Self simulated dataset

The SVM, logistic regression model and kNN algorithm used all 4 simulated biomarkers as explanatory variables to classify disease status. Following tuning of hyper-parameters for each SVM with corresponding Kernel function, it was found that the SVM with linear Kernel function and a cost value of *C* = 0.0125 achieved the best performance. It achieved 94% accuracy, with sensitivity of 79% and specificity of 100% and was significantly better than the NIR (P*<*0.001). The logistic regression model performed similarly, with accuracy 94%, sensitivity 86% and specificity 97%; it was also significantly better than the NIR (P*<*0.001). The SVM performed slightly better than the kNN algorithm, which achieved accuracy 90%, sensitivity 71% and specificity 97%; it was also significantly better than the NIR (P=0.002).

#### Case study of prostate cancer

The SVM, logistic regression model and kNN algorithm used the ratio of free to total PSA as the sole explanatory variable to classify disease status. Following tuning of hyper-parameters for each SVM, it was found that performance was comparable between the three SVM models; the SVM with linear kernel with cost value 0.0125 was used as it is computationally more efficient and the simplest to implement (less hyper-parameter choice). The SVM achieved an accuracy of 72%, sensitivity of 79%, specificity of 65% and the test against the NIR showed good significance (P=0.014). The logistic regression model performed similarly, with accuracy 72%, sensitivity 79%, specificity 65% and outperformed the NIR (P=0.014). The kNN algorithm slightly outperformed the SVM, by only a slight margin, with accuracy 75%, sensitivity 79% specificity 71%; the test against the NIR showed that it was significantly better (P=0.005).

The performance of the SVM can be assessed visually in Figure 10; as there was only one variable of interest in this classification, the ratio of free to total PSA is on the y-axis and a patient identifier is on the x-axis. The red shaded region represents the data points that would be classified as positive for prostate cancer by the SVM, and the green shaded region represents points that would be classified as negative. It can be seen that the SVM has a cut-off value of 0.2, a ratio value below this cut-off leads to the SVM classifying the patient as positive for prostate cancer. Actual data points from the validation dataset were then overlaid, represented by the darker red circles and darker green squares on the plot. The good accuracy achieved by the SVM can be seen, there were three patients that did not have prostate cancer that were wrongly classified as positive for prostate cancer (represented by green squares that lie in the red shaded region) and there were five patients that had prostate cancer that were wrongly classified as negative for prostate cancer (represented by red circles in the green shaded region).

**Figure (10).**
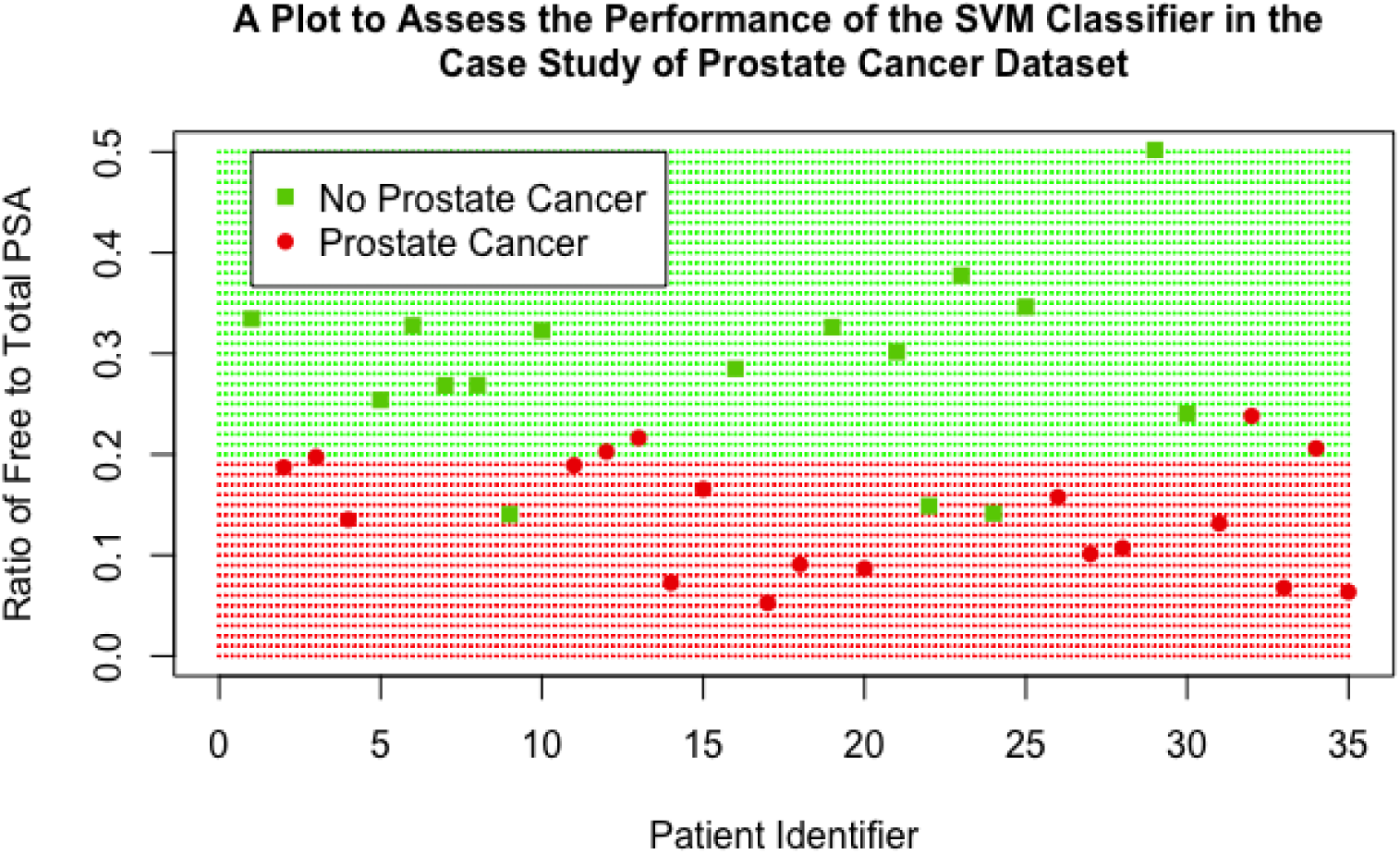
A scatter plot showing the performance of the final SVM classifier in the case study of prostate cancer dataset

#### Pancreatic cancer biomarkers

Two separate SVMs were fitted, one using y1 and y2 as explanatory variables and the other using log(y1) and log(y2) as explanatory variables, to assess the impact of transformations on the performance of the SVM; a logistic regression model and kNN algorithm were also implemented with log(y1) and log(y2) as explanatory variables. The tuning process showed that SVMs using linear and polynomial Kernel functions performed comparably for both the log and non-log case, so the linear was carried forward in both cases due to its computational efficiency and ease of implementation; the SVM using log had a cost value of 0.0625 and the SVM not using log had a cost value of 32. Performance between the 2 SVM models was comparable, showing that SVM performance was robust to transformations of the explanatory variables in this case; although their optimal specification was quite different, as displayed by their vastly different cost values. Performance measures were similar: accuracy (86% for log variables vs 86% for non-log variables), sensitivity (88% vs 80%) and specificity (0.82% vs 100%); both performed better than NIR (P=0.018 for both). Both SVMs slightly outperformed the logistic regression model, which achieved 81% accuracy, 88% sensitivity, 64% specificity and was not significantly different to the NIR (P=0.099). Both SVMs also outperformed the kNN algorithm, which achieved 78% accuracy, 84% sensitivity, 64% specificity and was not significantly different to the NIR (P=0.184).

The performance of the SVM (using log variables) can be assessed visually in 11, this plot displays log(y1) on the x-axis and log(y2) and the y-axis. The cut-off provided by the linear SVM can be clearly seen by observing the shaded background regions, the green shaded region represents the points that would be classified as negative for pancreatic cancer by the SVM and the red region represents points that would be classified as positive. Low values for both log(y1) and log(y2) lead to the SVM classifying the data as negative for pancreatic cancer, if a patient has either above a value of 5 then the SVM gives a positive classification. Real data points from the validation dataset were again overlaid, the high accuracy of the SVM is clear as only 5 patients were incorrectly classified (represented as a green square in the red region or a red circle in the green section).

#### Gene expression array

The SVM, logistic regression model and kNN algorithm used all 1,524 gene expression variables as explanatory variables. This caused the model tuning process to become computationally very expensive and challenging. In fact, the grid search in combination with cross-validation failed, R gave the error ‘Maximum number of iterations reached’. To circumvent this, a ‘sparse’ grid search was initially carried out; this involved performing a grid search over a reduced range i.e. *C* = {2^−5^, 1, 2^5^}, *γ* = {0.25, 1, 4} and *d* = {1, 2}, to narrow down where the best performance was. A full grid search in combination with cross-validation was then carried out over the following ranges, to find the optimal performance: *C* = {0.25, 0.5, 1, 2, 4}, *γ* = {0.25, 0.5, 1, 2, 4} and *d* = {1}. Following this extended tuning process, it was found the the SVM with linear Kernel and cost value 0.03125 performed the best; it achieved 86% accuracy, 83% sensitivity and 88% specificity; it was not significantly different to the NIR (P=0.086). The SVM out performed the logistic regression model, which achieved an accuracy of 64%, a sensitivity of 83% and a specificity of 50%; the logistic regression model was also not significantly different to the NIR (P=0.399). The kNN slightly outperformed the SVM in this case, with an accuracy of 93%, sensitivity 100% and specificity 88%; it was significantly better than the NIR (P=0.005). Although the kNN algorithm did perform better than the SVM, the effect was exaggerated due to the size of the testing dataset. In this case the testing set contained only 14 patients, so a small difference in the number of correct classifications between the two methods led to a large difference in accuracy. The SVM and kNN algorithm correctly classified 12 and 13 of the 14 patients respectively, the 7% increase in accuracy in favour of the kNN algorithm is therefore slightly misleading, the two methods in fact showed similar performance. A plot was not created for this dataset as there were 1,524 variables included for classification by the SVM and a plot would only show performance for two of these, this would not be representative of the overall performance of the SVM.

#### Risk reclassification (simulated)

The SVM, logistic regression model and kNN algorithm used all 3 simulated biomarker values as explanatory variables. As discussed, this dataset contained 10,000 observations, this caused the tuning process to become far too computationally expensive to implement a full hyper-parameter tuning process. Again, a sparse grid search was initially carried out in order to narrow down the search area; the sparse ranges were *C* = {2^−5^, 1, 2^5^}, *γ* = {0.25, 1, 4} and *d* = {1, 2}. Even with this reduced range, the tuning for the polynomial SVM failed after over an hour of carrying out the procedure, so the degree range was limited to being 1 only; calculating higher order polynomial functions is what causes the computational expense to increase, so restricting the degree hyper-parameter helped to limit computation time. After this sparse search, a full grid search in combination with cross-validation was then carried out over the following ranges: *C* = {0.25, 0.5, 1, 2, 4}, *γ* = {0.25, 0.5, 1, 2, 4} and *d* = 1. It was found that the SVM with radial Kernel with cost value 1 and *γ* =1 showed the best performance, it achieved an accuracy of 96%, sensitivity of 77% and a specificity of 98%. Both the logistic regression model and kNN algorithm performed similarly to the SVM: the LR achieved an accuracy of 96%, sensitivity 74% and specificity 98%; the kNN algorithm achieved an accuracy of 96%, sensitivity 78% and specificity 98%. All models were significantly different compared to the NIR (P*<*0.001 for all three).

The performance of the SVM can be assessed visually in Figure 12, this plot displays ‘x’ on the x-axis and ‘y’ and the y-axis; the variable ‘w’ was still included in the SVM classifier but is not represented in this plot. The cut-off region defined by the SVM can be clearly seen, the red shaded background region represents the data points that would be classified as positive for disease and the green region represents the points that would be classified as negative for disease; the interesting shape of this region was achieved by the use of the RBF Kernel. 25% of the data points from the validation set were overlaid on this plot to visually assess the performance of the SVM (including all data points was not feasible as they obscured the plot and no detail could be observed), the high accuracy is clear as only a minority of points were misclassified.

**Figure (11).**
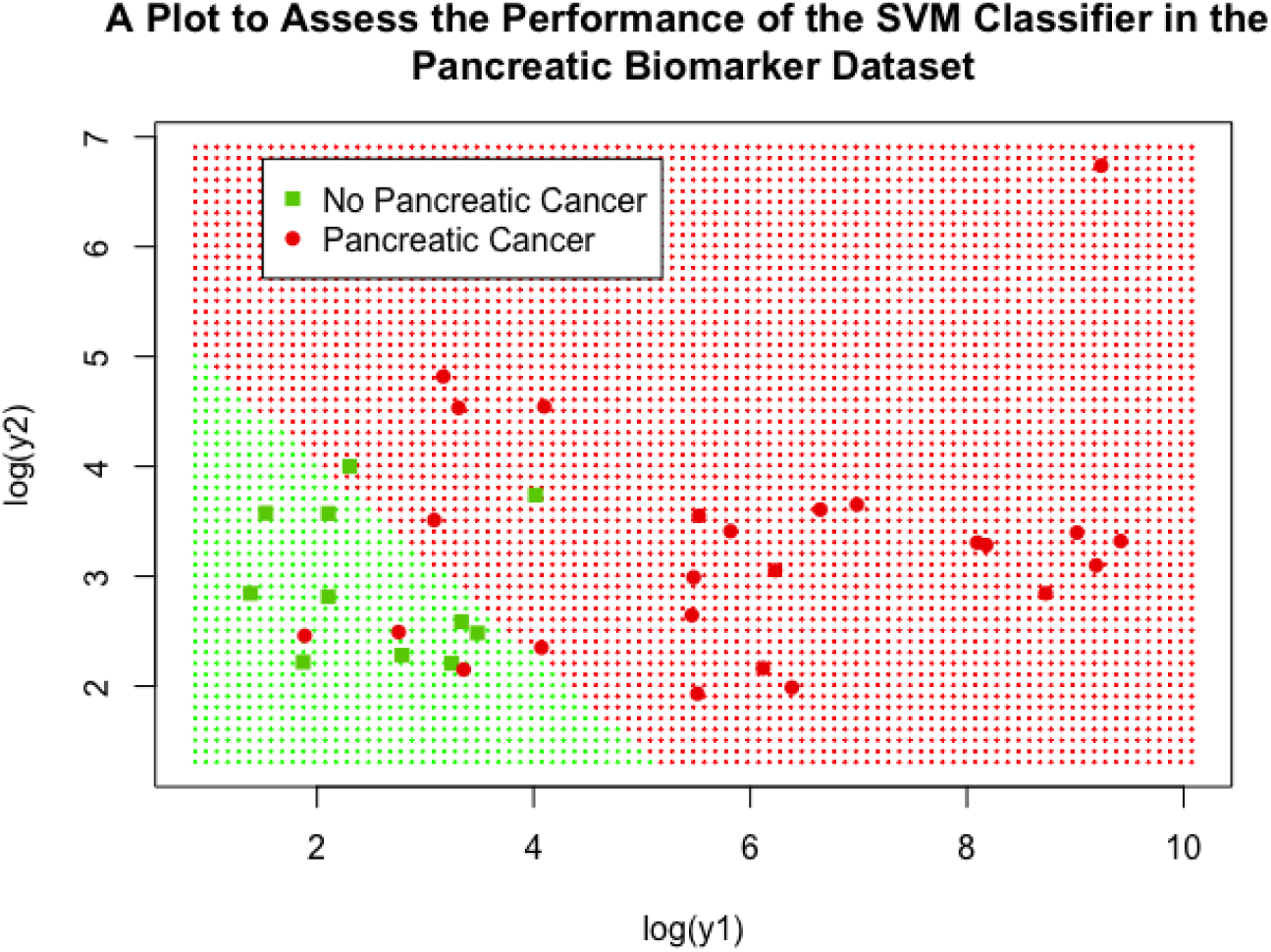
A scatter plot showing the performance of the final SVM classifier in the pancreatic biomarker dataset

**Figure (12).**
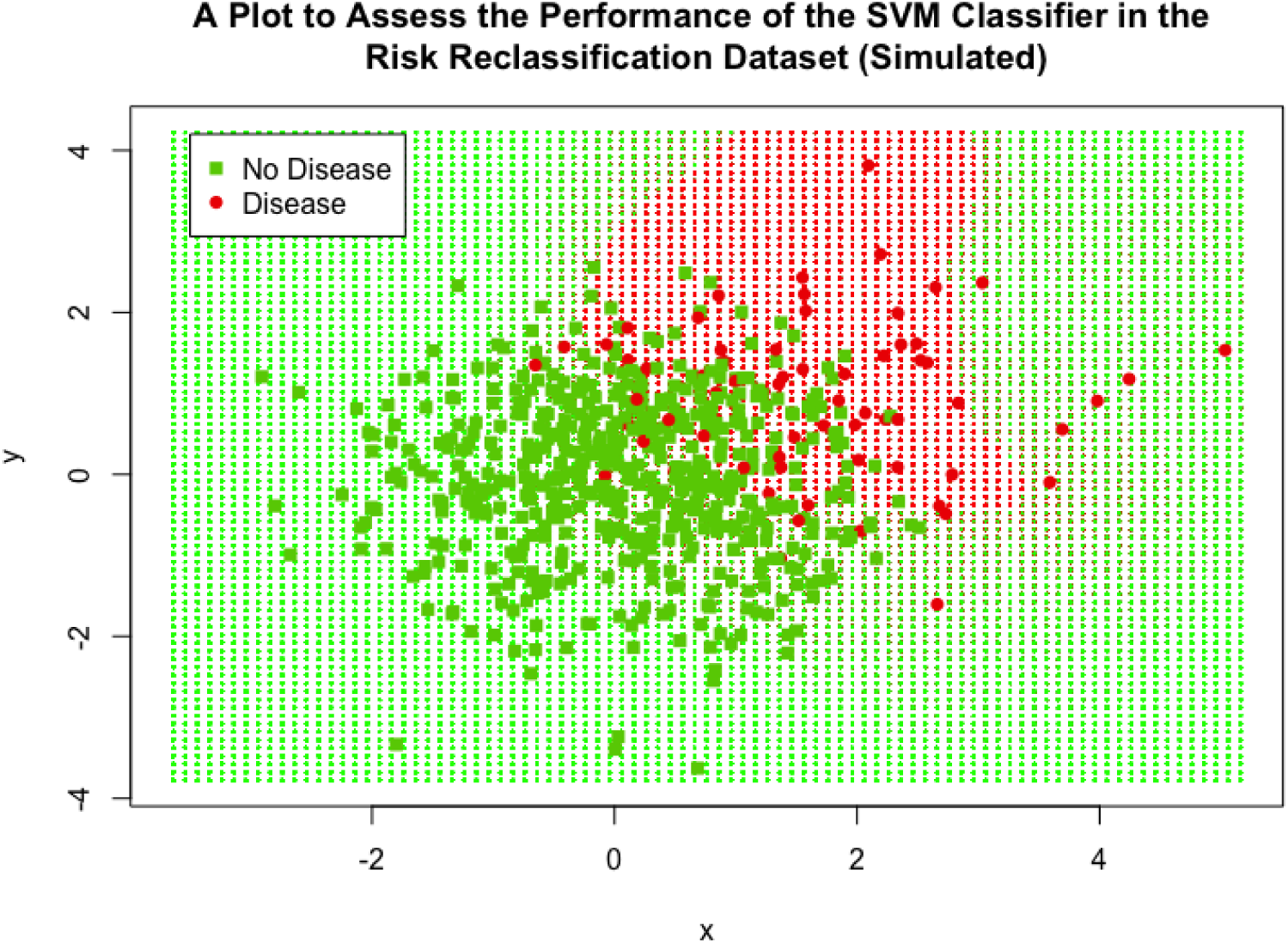
A scatter plot showing the performance of the final SVM classifier in the simulated risk reclassification dataset

## 5 Discussion

It has been demonstrated that SVMs outperform both a simple logistic regression model and an existing ML classification method (kNN) in a number of cases and show similar performance in others. Performance measures were comparable between the SVM, LR model and kNN algorithm in both small and large datasets (the case study of prostate cancer dataset and the risk reclassification dataset contained 141 and 10,000 observations respectively), displaying the wide applicability of SVMs to varying sizes of data. SVMs also showed robustness to data transformations, performance measures before and after application of a log transformation to the pancreatic cancer biomarker were similar; this is an attractive feature as many biological features often have skewed distributions. The SVM also outperformed the LR and kNN in this case. Many problems being faced in industry and research make use of complex, non-linear data, which can be easily incorporated by the SVM methodology, making them an appealing option in such cases. This was showcased when applying the SVM to gene expression data, the SVM showed superior performance measures when compared with the LR and performed similarly to the kNN.

A limitation of using SVMs is the dependency of the Kernel choice and its respective hyperparameters. The choice of which Kernel function to use is often subjective and data-dependent; generally, multiple Kernel functions are trialled and whichever achieves the best classification accuracy is then taken forward. Further, the tuning of hyper-parameters becomes computationally challenging when either the number of parameters or the number of observations becomes very large; this problem was encountered in both the gene expression dataset (*>*1500 variables) and the risk reclassification dataset (10,000 observations). Hsu, Chang and Lin [24] state that more advanced methods can be utilised over a grid search in conjunction with cross validation; many such algorithms for big data have been put forward [36]. This is, though, less of a concern for clinical studies since the sample size is usually in hundreds or at best a few thousand patients.

In conclusion, the wide applicability of SVMs for classification problems has been demonstrated, they show utility in complex problems and perform well in a variety of situations. They achieve optimal stratification of patient populations based on prognostic cancer biomarkers and are therefore an attractive option for stratification in oncology settings. In particular, the implementation of SVMs to quantitative prognostic biomarker datasets could play a valuable role

in observational studies by stratifying patient population into subgroups achieving differential clinical outcomes irrespective of treatment, or in classifying patients into risk categories, or indeed in using the biomarker signature as a stratification variable in a randomised clinical trial. Though outside the remit of this work, the SVM methodology and algorithms presented herein are ‘oven ready’ for implementation in predictive biomarkers by stratifying patient population into subgroups and identifying the one most likely to benefit from targeted cancer therapies.

